# Extensive post-transcriptional buffering of gene expression in the response to oxidative stress in baker’s yeast

**DOI:** 10.1101/501478

**Authors:** William R. Blevins, Teresa Tavella, Simone G. Moro, Bernat Blasco-Moreno, Adrià Closa-Mosquera, Juana Díez, Lucas B. Carey, M. Mar Albà

## Abstract

Cells responds to diverse stimuli by changing the levels of specific effector proteins. These changes are usually examined using high throughput RNA sequencing data (RNA-Seq); transcriptional regulation is generally assumed to directly influence protein abundances. However, the correlation between RNA-Seq and proteomics data is in general quite limited owing to differences in protein stability and translational regulation. Here we perform RNA-Seq, ribosome profiling and proteomics analyses in baker’s yeast cells grow in rich media and oxidative stress conditions to examine gene expression regulation at various levels. With the exception of a small set of genes involved in the maintenance of the redox state, which are regulated at the transcriptional level, modulation of protein expression is largely driven by changes in the relative ribosome density across conditions. The majority of shifts in mRNA abundance are compensated by changes in the opposite direction in the number of translating ribosomes and are predicted to result in no net change in protein level. We also identify a subset of mRNAs which is likely to undergo specific translational repression during stress and which includes cell cycle control genes. The study suggests that post-transcriptional buffering of gene expression may be more common than previously anticipated.

## Introduction

In recent years high throughput RNA sequencing (RNA-Seq) has become the method of choice for measuring shifts in gene expression between cells grown in different conditions \ However, diverse studies have shown that mRNA levels only partially explain protein levels in the cell^2–5^. In yeast, the correlation between mRNA and protein abundance is typically in the range 0.6-0.7^2^. In addition, the ratio between protein and mRNA levels may vary across different conditions^3^. For instance, substantial differences in this ratio have been observed during osmotic stress in yeast^6^ or after the treatment of human cells with epidermal growth factor^7^

In contrast to RNA-Seq, which measures the total amount of mRNA in the cell, ribosome profiling (Ribo-Seq) only captures those mRNAs that are being actively translated^8^. Each Ribo-Seq read corresponds to one translating ribosome, providing a quantitative view of the amount of protein produced by the cell at any given time. Although this remains an indirect estimate of protein abundance, it has several advantages over proteomics, such as the fact that with Ribo-Seq virtually all translated sequences can be captured, and that one can apply the same pipelines and statistical methods as for RNA-Seq to identify differentially expressed genes.

The response to oxidative stress in the yeast *Saccharomyces cerevisiae* involves a general decrease in mRNA translation initiation as well as the selective transcriptional activation of a set of proteins involved in the maintenance of the redox state of the cell^9–11^. A previous study reported changes in the ratio between the normalized number of Ribo-Seq and RNA-Seq reads, or translational efficiency (TE), of hundreds of genes upon oxidative stress^9^, suggesting extensive translational regulation. However, changes in TE alone do not necessarily imply changes in the abundance of the translated proteins. Here, by performing a separate analysis of Ribo-Seq and RNA-Seq data, we show that the majority of genes that show statistically significant differences at the RNA-Seq level do not show similar differences at the Ribo-Seq level, suggesting that, in most cases, changes in mRNA abundance are compensated by changes in ribosome density and are not propagated to the protein level. Our approach also uncovers a subset of differentially expressed genes in which regulation appears to be mainly exerted at the translational level.

## Results

### Ribosome profiling experiments in normal and stress conditions

We extracted ribosome-protected RNA fragments, as well as complete polyadenylated RNAs, from *Saccharomyces cerevisiae* grown in rich media (normal) and in H_2_O_2_-induced oxidative stress conditions (stress)(Figure 1). We then sequenced the ribosome-protected RNA fragments (Ribo-Seq) as well as complete mRNAs (RNA-Seq) using a strand-specific protocol. The Ribo-Seq data provided a snapshot of the translatome, each read corresponding to one translating ribosome, whereas the number of RNA-Seq reads mapping to a gene was used to quantify the relative abundance of the transcript.

**Figure 1.**
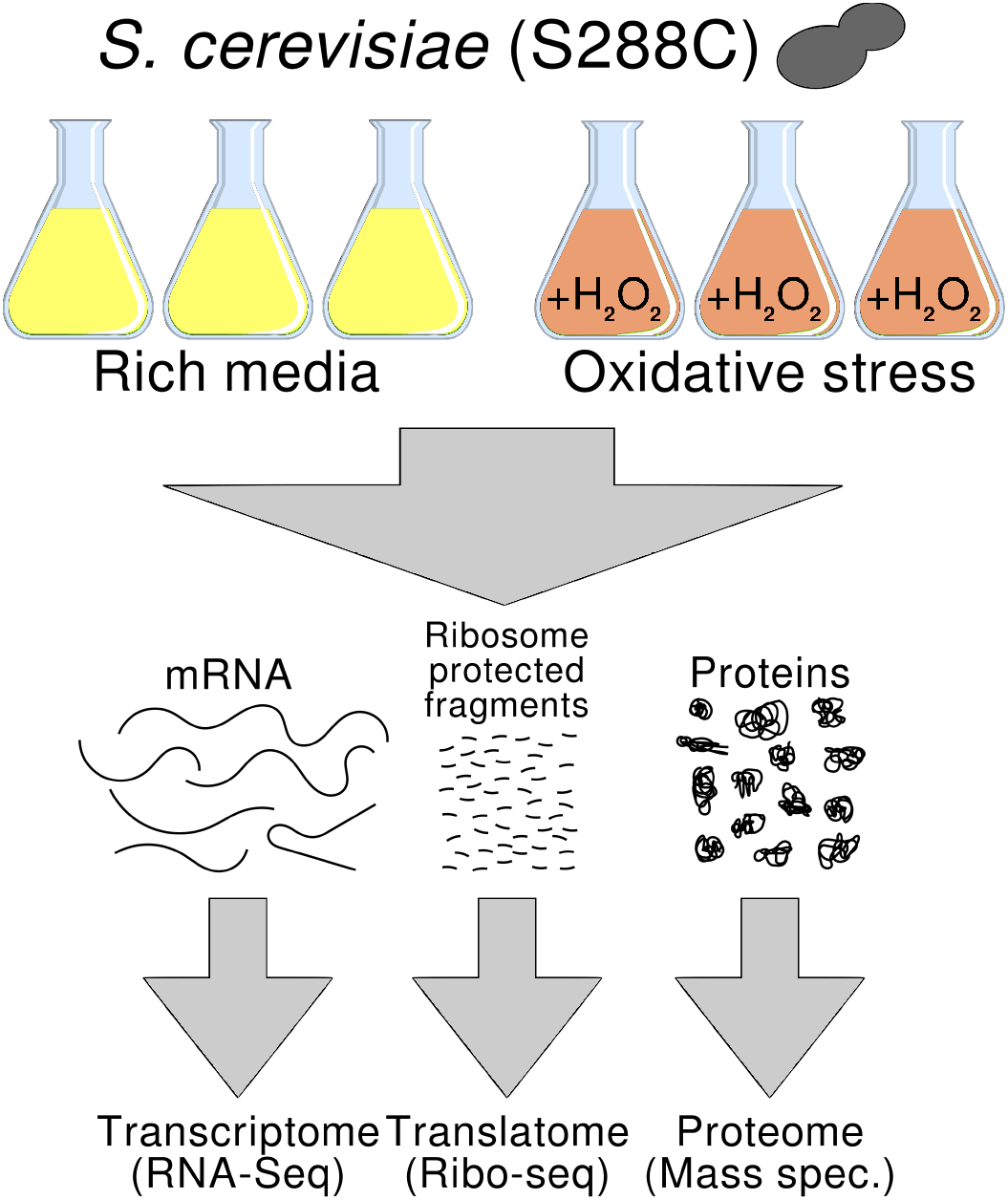
Experimental design. Baker’s yeast (S. *cerevisiae*) was grown in rich media and oxidative stress conditions in parallel. The cultures were used to extract total RNA, ribosome-protected RNA fragments and proteins.

After quality control of the sequencing reads we obtained 31-36 million Ribo-Seq reads and 12-15 million RNA-Seq reads per sample (Supplementary Table S1). We mapped the reads to the genome and generated a table of read counts per gene for each of the samples. After filtering out non-expressed genes (see Methods), the table contained data for 5,419 *S. cerevisiae* annotated genes (ORFs).

We normalized the RNA-Seq and Ribo-Seq table of counts by calculating normalized counts per million (CPM) in logarithmic scale, or log_2_CPM (Supplementary Figure S1). The correlation coefficient between the average Ribo-Seq and RNA-Seq log_2_CPM expression values was 0.84 in normal conditions and 0.87 in stress conditions (Figure 2 A and B, respectively). As the differences in log_2_CPM between RNA-Seq or Ribo-Seq replicates were negligible (Figure 2 C and D, Supplementary Table S2), these values reflect the amount of disagreement between total mRNA and translated protein abundances.

**Figure 2.**
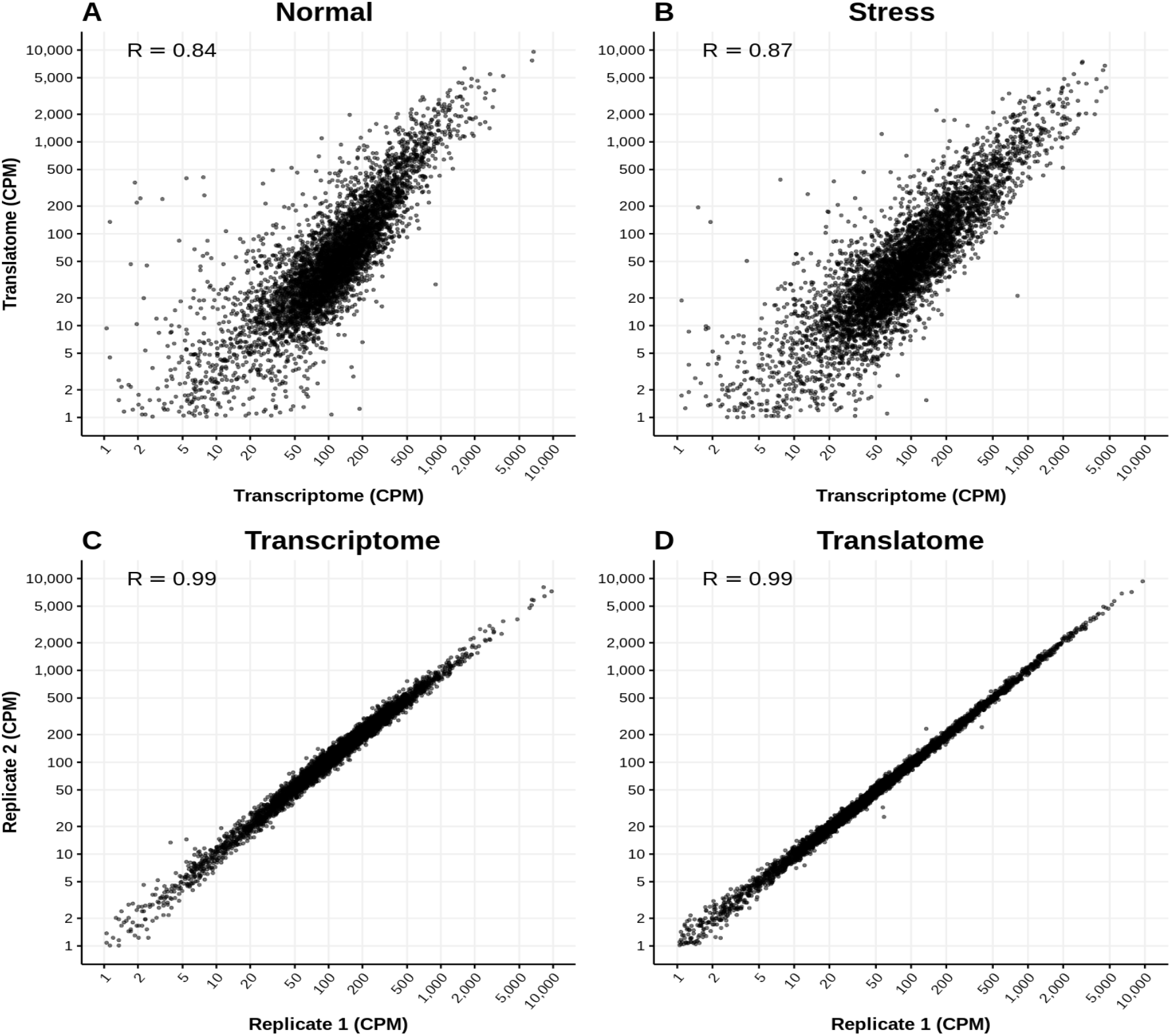
Representative gene expression correlations between RNA sequencing samples. **A**. RNA-Seq normal replicate 1 *versus* Ribo-Seq normal replicate 1. **B**. RNA-Seq stress replicate 1 *versus* Ribo-Seq stress replicate 1. **C**. RNA-Seq normal replicate 1 *versus* RNA-Seq normal replicate 2. **D**. Ribo-Seq normal replicate 1 *versus* Ribo-Seq normal replicate 2. Expression units are CPM in logarithm scale; R: Spearman correlation value. N: normal growth conditions (two replicates N1 and N2); S: stress conditions (two replicates S1 and S2).

### Ribo-Seq shows a higher correlation with proteomics than RNA-Seq

The next step was to compare the quantification of gene expression by RNA-Seq and Ribo-Seq to that obtained using proteomics. We extracted the protein fraction from yeast grown in normal and stress conditions and estimated the abundance of different yeast proteins, i.e. the proteome, using mass spectrometry information (Figure 1). We could reliably quantify the protein products of 2,200 genes (see Methods), representing about 40% of the genes quantified by RNA-Seq or Ribo-Seq. Normalized protein abundances between pairs of proteomics replicates showed correlation coefficients in the range 0.83-0.93 (Supplementary Table S3), lower than for RNA-Seq or Ribo-Seq replicates (>0.99).

In normal conditions the correlation coefficient between the transcriptome (RNA-Seq) and the proteome relative abundance units was 0.46. This increased to 0.71 when comparing the translatome (Ribo-Seq) and the proteome units (Figure 3). This indicates that Ribo-Seq-based quantification of gene expression provides a more accurate picture of protein abundance than RNA-Seq data. The average correlation coefficient between the three pairs of proteome replicates was 0.91, setting up a maximum value for any correlation. Differences between RNA-Seq and proteomics quantification estimates may arise because of differences in the half life of the proteins with respect to their cognate mRNAs as well as variations in the translation rate or ribosome density across the transcripts. As the value of 0.71 (Ribo-Seq *versus* proteomics) is intermediate between 0.46 (RNA-Seq *versus* proteomics) and 0.91 (proteomics replicates), the two above mentioned factors appear to be relevant to explain the strong uncoupling between mRNA and protein abundance in this system.

**Figure 3.**
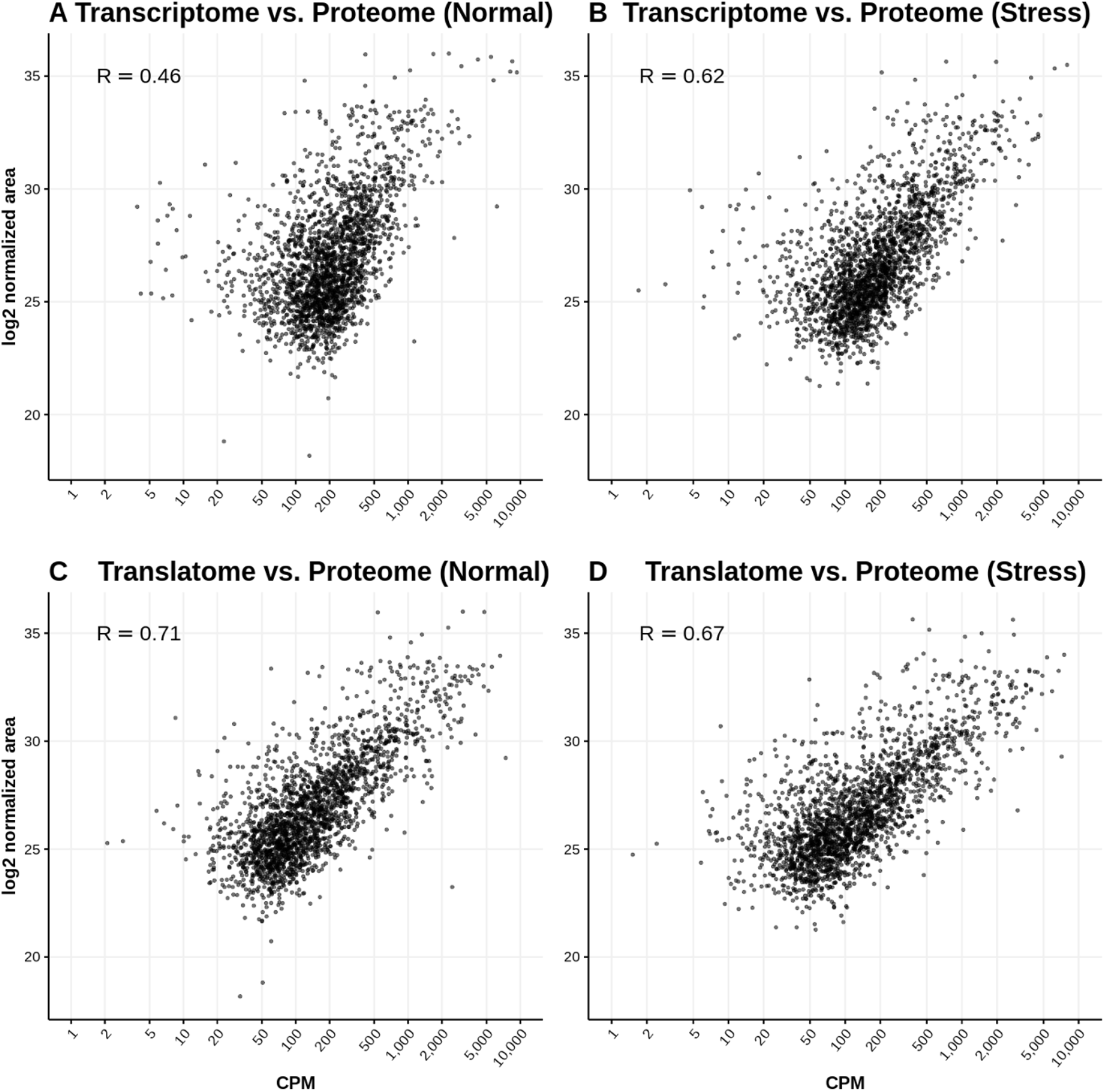
Proteomics shows a stronger correlation with Ribo-Seq than with RNA-Seq data. **A**. RNA-Seq *versus* proteomics, normal growth conditions. **B**. RNA-Seq *versus* proteomics, oxidative stress. **C**. Ribo-Seq *versus* proteomics, normal growth conditions. **D**. Ribo-Seq *versus* proteomics, oxidative stress. CPM: counts per million for RNA-Seq and RNA-Seq data (represented in logarithmic scale, average between replicates). log_2_ normalized area: relative abundance for proteomics data (average between replicates). R: Spearman correlation value. Plot and correlations comprise 2200 genes for which ≥3 unique peptides were detected by LCMSMS.

In stress conditions the correlation coefficient between the transcriptome and proteome was 0.62, somewhat higher than in normal conditions. The correlation coefficient between the translatome and the proteome was 0.67, again higher than the same value between the transcriptome and the proteome but lower than the correlation between the proteome stress replicates (0.86). Taken together, these results are consistent with the hypothesis that differences in ribosome density play a role in modulating protein expression.

### Analysis of three nucleotide periodicity

In actively translated regions mapped Ribo-Seq reads exhibit a characteristic three nucleotide periodicity that results from the codon-to-codon ratcheting movement of the ribosome along the coding sequence^8^. We used the program RibORF^10^ to assess the nucleotide periodicity and homogeneity of the Ribo-Seq reads in the annotated coding sequences. According to this analysis, in the vast majority of genes (98%, 5198 out of 5304 analyzed genes) the annotated ORF appeared to be translated in both normal and stress conditions, validating our approach of considering all the reads that mapped to the annotated ORFs for the quantification of protein translation.

In a small fraction of genes, however, we found evidence of alternative translated ORFs (Supplementary Table S4). One example was TOS8, which encodes a homeodomain-containing transcription factor. In this gene active translation of the canonical 831 amino acid long protein by RibORF was only detected in stress conditions; in contrast, a protein of only 81 amino acids was the main translated polypeptide in rich media. The shorter alternative ORF was on a different reading frame to the main protein product and showed no homology to any previously characterized protein. These cases illustrate how detailed examination of the distribution of the Ribo-Seq reads may help uncover proteins that have remained hidden within longer ORFs.

### Ribo-Seq estimates of changes in gene expression are more conservative

We next calculated the gene expression level fold change (FC) between the two conditions, using RNA-Seq and Ribo-Seq data separately. The log_2_FC distribution based on the Ribo-Seq data had a lower variance than the log_2_FC distribution using RNA-Seq data (Figure 4A). This indicated a higher range of variation in the mRNA levels, as estimated by RNA-Seq, than in the ribosome-protected fragments. This was consistent with the existence of post-transcriptional buffering of gene expression, as also reported for inter-specific gene expression comparisons of *S. cerevisiae* and *S.paradoxus*^11^.

**Figure 4.**
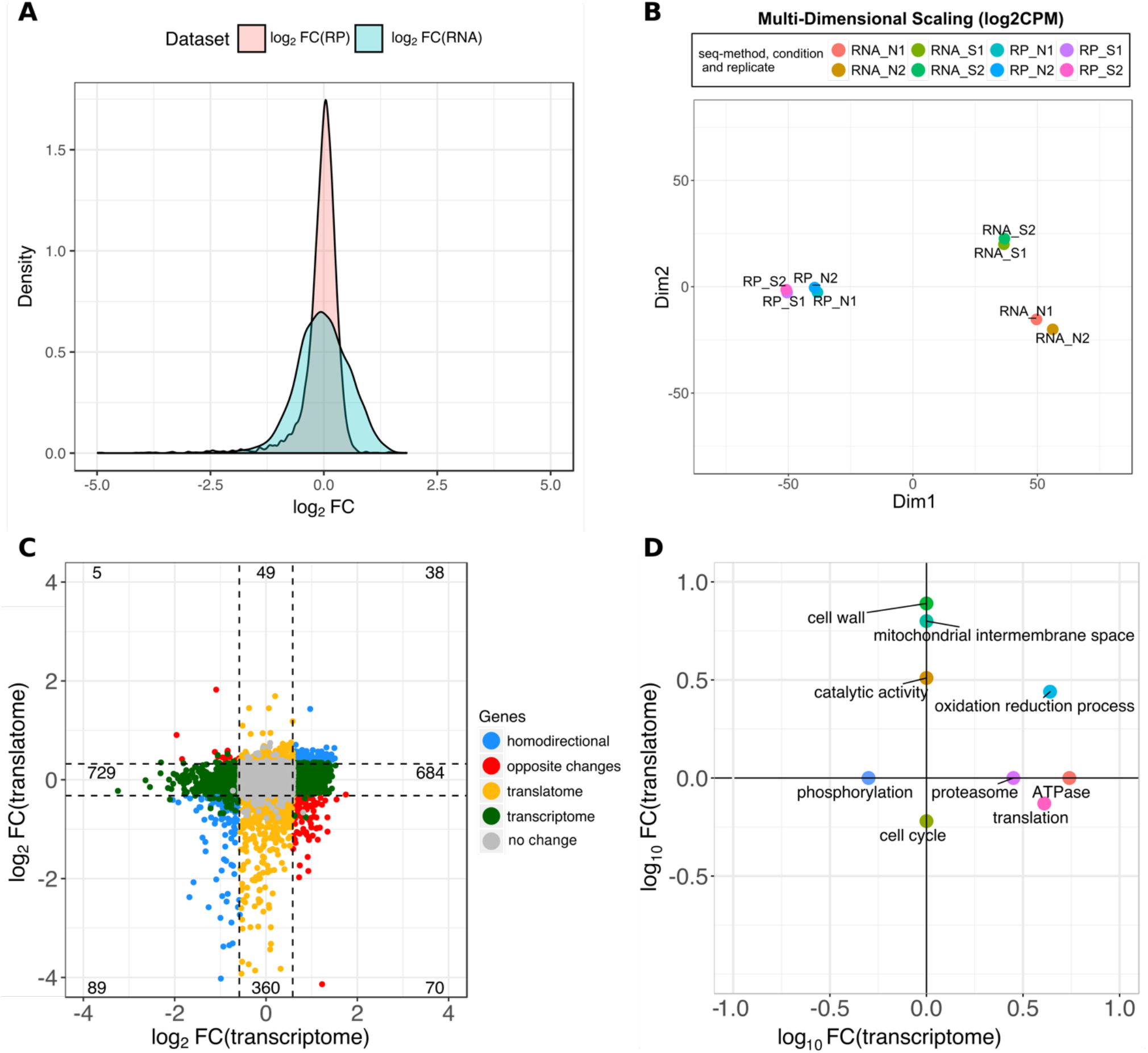
Integrated analysis of RNA sequencing and ribosome profiling data. **A**. Distribution of gene expression fold change (FC) values. FC was calculated as the ratio between the number of reads in oxidative stress and normal conditions. We took the average number of reads per gene among the replicates. The standard deviation of log_2_FC was 0.44 for Ribo-Seq (RP) and 0.57 for RNA-Seq (RNA). **B**. Multidimensional scaling (MDS) plot using the gene expression values of each sample. MDS was based on the log_2_CPM values for each gene. Data was for 5,419 S. cerevisiae genes. RP: Ribo-Seq data; RNA: RNA-Seq data; N: normal growth conditions; S: stress conditions. Two sequencing replicates were generated per condition. **C**. Correlation between log fold change (FC) gene expression values. The X axis corresponds to the RNA-Seq data, or transcriptome, the Y axis to the Ribo-Seq data, or translatome. Coloured dots correspond to differentially expressed genes. In the legend homodirectional means up-regulated, or down-regulated, both at the transcriptome and translatome levels; opposite_change is up-regulated at one level and down-regulated at the other one; translatome means significant differences in Ribo-Seq only; transcriptome means significant differences in RNA-Seq only. **D**. Significant gene functional classes among differentially expressed genes. Shown is a 2-D plot of the enrichment score values, in logarithmic scale, provided by the software DAVID for differentially expressed genes using RNA-Seq (transcriptome) or Ribo-Seq (translatome) data. Significant enrichment scores are associated with a p-val < 0.05. Functional classes associated with positive values are significantly enriched among up-regulated genes, and functional classes with negative values are significantly enriched among down-regulated genes. Non-significant enrichment scores are given a value of 0 in the plot.

We considered the possibility that the about 2.5 times higher number of Ribo-Seq reads than RNA-Seq read in the original datasets biased the comparison of log_2_FC distributions. In order to test it we subsampled the mapped reads so as to have a similar number of reads in all the RNA-Seq and Ribo-Seq samples (Supplementary Tables S5 and S6). The results were very similar to those observed without subsampling (Supplementary Figure S2), indicating that these observations have a biological origin.

We also used an alternative method, multidimensional scaling (MDS)^12^, to quantify the distance between Ribo-Seq and RNA-Seq gene expression measurements (Figure 4B). We found that the distance between Ribo-Seq normal and stress conditions was shorter that the distance between RNA-Seq normal and stress conditions, which was consistent with the previous observation that log_2_FC variance was lower for Ribo-Seq than for RNA-Seq.

### Extensive post-transcriptional buffering of gene expression

We next performed differential gene expression analysis, separately for Ribo-Seq and RNA-Seq data, using multivariable linear regression with the Limma package^13^. Limma provides a list of differentially expressed genes with the corresponding adjusted p-values. We selected genes with an adjusted p-value < 0.05 and a log_2_FC larger than one standard deviation; the latter corresponded to a minimum FC of 1.49 for RNA-Seq data and 1.36 for Ribo-Seq data. We used the standard deviation instead of a fixed value to accommodate for the differences in the width of the log_2_FC distributions. The number of genes that were differentially expressed was 1,530 for RNA-Seq and 536 for Ribo-Seq.

The correlation between RNA-Seq and Ribo-Seq gene log_2_FC values was quite low (0.18), indicating an important disconnect between the two kinds of data (Figure 4C). Only 127 genes showed a significant change in the same direction i.e. homodirectional changes. Genes that were up-regulated during stress according to both RNA-Seq and Ribo-Seq included protein functions known to be activated at the transcriptional level in response to stress, such as hexoquinases or heat shock proteins^14^ The number of genes annotated with the Gene Ontology (GO) term ‘oxidation reduction process’ was similar for RNA-Seq or Ribo-Seq up-regulated genes (17 and 15, respectively), supporting that these genes are essentially regulated at the level of transcription and can be effectively detected with both kinds of sequencing data.

The vast majority of genes were only significant at the transcriptome or the translatome levels (1,413 and 409 genes, respectively; Figure 4C). The first group was formed by genes that showed significant changes in relative transcript abundance but not in the relative number of ribosome-protected fragments, supporting extensive post-transcriptional buffering of gene expression. The data indicated that about a quarter of the genes in the genome may be undergoing compensatory changes: when mRNA levels increase ribosome density per transcript decreases and the other way round. The levels of the proteins encoded by these genes are not expected to change despite significant changes in the corresponding mRNA abundance.

The second group, translatome-only differentially expressed genes, represented cases in which mRNA levels did not change but the density of ribosomes per transcript showed a significant increase or decrease in stress relative to normal. This would be consistent with the expression of these genes being primarily modulated at the level of translation. We identified many more genes under differential translational repression than activation (360 *versus* 49, Figure 4C), suggesting that the former mechanism may be more prevalent that the first one in response to stress.

Finally, we found a subset of cases showing opposite changes in RNA-Seq and Ribo-Seq data. The main group was formed by 70 genes showing increased mRNA levels but decreased translation in stress *versus* normal. One simple explanation would be that, for these genes, there is an mRNA fraction that is stored in a translational inactive highly stable form, whereas the rest is translated at the usual level. More complex scenarios could involve a combination of transcriptional and translational regulatory events.

### Dissecting differential regulation by functional class

To better understand the biological relevance of our observations, we investigated if certain functional classes were significantly enriched among the sets of differentially expressed genes. We used DAVID^15^ to identify significantly over-represented functional clusters (Figure 4D). Only one class, ‘oxidation-reduction process’, was enriched among genes up-regulated during stress both using RNA-Seq and Ribo-Seq data. This is consistent with transcriptional activation of this set of genes upon stress, increasing the signal for both total mRNA and the translated fraction.

Three other classes – ‘translation’, ‘ATPase’ and ‘proteasome’ – showed increased mRNA levels during stress, but this was not reflected in an increase in the translated fraction. These classes may be particularly prone to undergo compensated mRNA changes. Among genes that were differentially expressed only when we used Ribo-Seq data ‘cell wall’, ‘mitochondrial intermembrane space’ and ‘catalytic activity’ were enriched among up-regulated genes, whereas ‘cell cycle’ was enriched among down-regulated genes (Figure 4D).

### Translational efficiency and protein level changes

To obtain further insights into the regulatory mechanisms of gene expression during oxidative stress in yeast we also compared the translational efficiency (TE; Ribo-Seq normalized counts divided by RNA-Seq normalized counts) of the different genes in the two conditions using the program Ribodiff^16^. We detected 470 genes that showed significantly increased TE during stress (adjusted p-value < 0.05; see Methods); about 82% of them were cases in which the relative mRNA levels had decreased during stress but this change had been compensated by an increase in ribosome density so that no significant changes in the amount of translated protein would be expected (transcriptome downregulated, Table 1). In only about 3% of cases increased TE was associated with translational activation and increased protein production (translatome upregulated, Table 1).

**Table 1.** Genes with significantly increased or decrease translational efficiency during oxidative stress. TE: gene translational efficiency. Ribodiff p-value < 0.05 and | log_2_(TE_stress_/TE_normal_)| > *0.67.* Translatome/Transcriptome defintions as in Figure 5.

|  | Translatome<br>upregulated | Translatome<br>downregualted | Transcriptome<br>upregulated | Transcriptome<br>downregulated | Other |
| --- | --- | --- | --- | --- | --- |
| Increased<br>TE under<br>stress | 14 | 0 | 0 | 385 | 71 |
| Decreased<br>TE under<br>stress | 0 | 208 | 356 | 0 | 150 |

In the case of genes with significantly lower TE in stress than in normal conditions the percentage of compensatory cases was also the predominant scenario, accounting for 50% of the genes in the class (356 out of 714, Table 1). The second most numerous group were genes likely to be actively repressed at the level of translation, accounting for 29% of the genes with significantly decreased TE (29%). The latter genes showed no change in mRNA levels but the relative number of associated ribosomes was lower in stress than in normal conditions, which would be expected to lead to a decrease in the protein levels. This group included 12 genes from the cell cycle functional category (Supplementary Table S7).

## Discussion

The adaptation of organisms to variations in different environmental conditions is associated with the activation or repression of gene expression. These changes are usually studied at the level of complete mRNA molecules using microarrays or next generation sequencing. However, changes in mRNA concentration do not necessarily reflect changes in their encoded protein products^7,11^.

Here we have explored the usefulness of ribosome profiling data to close the gap between mRNA and protein abundance estimates. Each ribosome profiling read corresponds to one translating ribosome and thus the number of reads that map to a gene reflects the amount of protein that is being made^8,17^ Numerous recent studies have used ribosome profiling to gain insights into novel translation regulatory mechanisms^18,19^ or to discover new translated RNA sequences^20–23^. However, there is a lack of studies addressing how ribosome profiling can be used to improve the estimates of protein abundance changes over RNA-Seq-based estimates. Our study shows that Ribo-Seq provides better estimates of protein abundance than RNA-Seq and that the results of differential gene expression analyses are drastically altered if we use Ribo-Seq of RNA-Seq as the source sequencing data.

The abundance of the different proteins in the cells is usually estimated using mass spectrometry proteomics data^24,25^. This provides a direct measurement of protein abundance that can account for the variations in the stability of different proteins; however, proteomics methods are much less sensitive than current RNA sequencing approaches and not all proteins can be detected in routine analyses^26^. In addition, the results obtained with high-throughput sequencing are more reproducible across biological replicates than those obtained with mass spec proteomics; this confers the former studies increased power to perform differential gene expression analyses.

Previous studies in yeast indicated that Ribo-Seq showed a higher correlation with proteomics data than RNA-Seq, but these conclusions were drawn after comparing data obtained from different laboratories^8^. Here we generated RNA-Seq, Ribo-Seq and proteomics data for yeast grown in identical conditions, leading to less biased comparisons. These results support the hypothesis that the Ribo-Seq read counts provide a better approximation to protein levels than RNA-Seq read counts.

We observed that many of the genes that were detected as significantly up- or down-regulated in stress by RNA-Seq did not show any significant changes using the Ribo-Seq data, indicating frequent post-transcriptional buffering of gene expression. Intriguingly, studies comparing the expression of orthologous genes from closely related species have also reported that gene expression is in general more variable when measured by RNA-Seq than Ribo-Seq^11,27^ We found that, during oxidative stress, genes encoding ribosomal proteins and members of the proteasome and ATPase complexes tended to show increased mRNA levels but, at the same time, the rate of translation decreased. We also have to consider that some mRNAs could be transiently stored in P-bodies or stress granules^28–30^, becoming inaccessible to the translation machinery. Translation of these transcript could be rapidly reactivated when the stress disappears.

Transcripts encoding proteins involved in the cell cycle appeared to be modulated differently. In this case there was no apparent change in the number of mRNA molecules but ribosome density decreased, presumably reflecting lower translation rates. Repression of this class of proteins may be related to a slow down of cell division under stress; the cells grown under oxidative stress showed approximately half doubling times when compared to those grown in rich media.

The results of this study illustrate the importance of performing ribosome profiling experiments to differentiate between changes in mRNA that are likely to result in changes in the protein levels to those that are not. Although obtaining Ribo-Seq data is more labour-intensive than RNA-Seq, the protocols are being simplified and its use is rapidly growing^31–33^. The methodological framework we have developed can be applied to other datasets and help advance our understanding of gene regulation in other conditions.

## Methods

### Biological material

We grew *S. cerevisiae* (S288C) in 500 ml of rich media^34^. In order to induce oxidative stress, 30 minutes before harvesting we added diluted H_2_O_2_ to the media for a final concentration of 1.5 mM. The cells were harvested in log growth phase (OD600 of ~0.25) via vacuum filtration and frozen with liquid nitrogen. !

### Ribosome profiling

In order to capture ribosome protected mRNAs, cyclohexamide was added one minute before the cells were harvested. Cyclohexamide is commonly used as a protein synthesis inhibitor in order to prevent ribosome run-off and the subsequent loss of ribosome-transcript complexes. One third of each culture was used for ribosome profiling (Ribo-Seq); the rest was reserved for RNA-Seq.

Cells were lysed using the freezer/mill method (SPEX SamplePrep); after preliminary preparations, lysates were treated with RNaseI (Ambion), and subsequently with SUPERaseIn (Ambion). Monosomal fractions were collected; SDS was added to stop any possible RNAse activity, then samples were flash-frozen with N2(l). Digested extracts were loaded in 7%-47% sucrose gradients. RNA was isolated from monosomal fractions using the hot acid phenol method. Ribosome-Protected Fragments (RPFs) were selected by isolating RNA fragments of 28-32 nucleotides (nt) using gel electrophoresis. The preparation of sequencing libraries for Ribo-Seq and RNA-Seq was based on a previously described protocol^35^. Pair-end sequencing reads of size 35 nucleotides (2×35bp) were produced for Ribo-Seq and RNA-Seq on MiSeq and NextSeq platforms, respectively. The data has been deposited at NCBI Bioproject PRJNA435567 (https://www.ncbi.nlm.nih.gov/bioproject/435567).

### Processing of the sequencing data

The RNA-Seq data was filtered using Trimmomatic with default parameters (version 0. 36)^36^. In the Ribo-Seq data we discarded the second read pair as it was redundant and of poorer quality than the first read, and then used Cutadapt^37^ to eliminate the adapters and to trim five and four nucleotides at 5’ and 3’ edges, respectively. Ribosomal RNA was depleted from the Ribo-Seq data *in silico* by removing all reads which mapped to annotated rRNAs. Ribo-Seq reads shorter than 25 nucleotides were not used.

After quality check and read trimming, the reads were aligned against the *S. cerevisiae* genome (S288C R64-2-1) using Bowtie 2^38^. For annotation we used a previously generated *S. cerevisiae* transcriptome containing 6,184 annotated coding sequences plus 1,009 non-annotated assembled transcripts (see Supplementary data). SAMtools^39^ was used to filter out unmapped reads.

We counted the number of reads that mapped to each gene with HTSeq-count^40^. We used the mode ‘intersection strict’ to generate a table of counts from the data; the procedure removed about 5% of the reads in the case of RNA-Seq, and 8% in the case of Ribo-Seq. Only genes in which the average read count of the two replicates was larger than 10 in all conditions (normal and stress, for RNA-Seq and for Ribo-Seq) were kept. The filtered table of counts contained data for 5,419 genes; nearly all of them corresponded to annotated genes (5,312 genes).

For subsampling the number of mapped reads we used SAMtools^39^. We used the function ‘samtools view’ with option ‘-s 0.X’, where X is the percentage of reads that we wish to keep.

### Analysis of three nucleotide periodicity in the mapped Ribo-Seq reads

We used RibORF^10^ to analyze the mapped Ribo-Seq. We analyzed all possible ORFs with a minium length of 9 amino acids and at least 10 mapped reads. WE analyzed 5,304 annotated ORFs. RibORF counts the number of reads that fall in each frame and calculates the distribution of reads along the length of the ORF. We used the original proposed cutoff (score > 0.7) to predict translated ORFs.

### Quantification of protein abundance by mass spectrometry

For our proteomics experiment, we analysed 3 replicates per condition by LCMSMS using a 90-min gradient in the Orbitrap Fusion Lumos. These samples were not treated with cyclohexamide. As a quality control measure, BSA controls were digested in parallel and ran between each sample to avoid carry-over and assess the instrument performance. The peptides were searched against SwissProt Yeast database, using the Mascot v2.5.1 search algorithm. The search was performed with the following parameters: peptide mass tolerance MS1 7 ppm and peptide mass tolerance MS2 0.5 Da; three maximum missed cleavages; trypsin digestion after K or R except KP or KR; dynamic modifications oxidation (M) and acetyl (N-term), static modification carbamidomethyl (C). Protein areas were obtained from the average area of the three most intense unique peptides per protein group. Considering the data from all 6 samples, we detected proteins from 3,336 genes. We limited our quantitative analysis to a subset of 2,200 proteins which had proteomics hits for at least 3 unique peptides; this filter eliminates noise arising from technical challenges of quantifying lowly abundant proteins with LCMSMS.

### Differential gene expression analysis

The table of counts was normalized to log_2_ Counts per Million (log_2_CPM) using the function ‘cpm’ in the R package edgeR^41^. Before performing differential gene expression analysis, we normalized the data using Trimmed Mean of M-values (TMM) from the same package. Finally, we applied the Limma voom method^13^ to identify differentially expressed genes, separately for RNA-Seq and Ribo-Seq data (adjusted p-value < 0.05 and |log_2_FC| > 1 SD(log_2_FC)).

We applied the same pipeline to the proteomics data using normalized area values as a quantitative measure of protein abundance. To ensure robustness of the differential expression analysis we used genes which had at least 3 unique peptides and could be quantified in all 6 replicates (1,580 genes); the procedure did not identify any significantly up or down regulated genes, using an adjusted p-value < 0.05. Low sensitivity of this procedure is expected considering the relatively poor correlation of the mass spec replicates (r between 0.83 and 0.93).

### Analysis of functional clusters

We identified significantly enriched functional clusters in differentially expressed genes using DAVID^15^. The analysis was done separately for over- and under-expressed genes and for RNA-Seq and Ribo-Seq derived data. Only clusters with enrichment score ≥ 1.5 and adjusted p-val < 0.05 were retained. In each cluster we chose a representative Gene Ontology (GO) term^42^, with the highest number of genes inside the cluster. Figure 4 integrates the results obtained with the Ribo-Seq and the RNA-Seq data, the log_10_ fold enrichment of the significant GO terms is plotted.

### Analysis of translational efficiency

We searched for genes with significantly increased or decreased translational efficiency (TE)^8^ using the RiboDiff program^16^. We selected genes significant at an adjusted p-value < 0.05 and showing log_2_(TE_stress_/TE_normal_) higher than 0.67 or lower than −0.67 (plus or minus one standard deviation of the distribution).

## Acknowledgements

We acknowledge the Proteomics Unit of Center for Regulatory Genomics and Universitat Pompeu Fabra for the isolation of proteins from yeast cultures. We are also grateful to Robert Castelo for advice during this project. The work was funded by grants BFU2015-65235-P, BFU2015-68351-P and BFU2016-80039-R, from Ministerio de Economía e Innovación (Spanish Government) – FEDER (EU), and from grant PT17/0009/0014 from Instituto de Salud Carlos III – FEDER. We also received funding from the “Maria de Maeztu” Programme for Units of Excellence in R&D (MDM-2014-0370) and from Agència de Gestió d’Ajuts Universitaris i de Recerca Generalitat de Catalunya (AGAUR), grant number 2014SGR1121, 2014SGR0974, 2017SGR01020 and, predoctoral fellowship (FI) to W.B. We also acknowledge support from the EU Erasmus Programme to T.T.

## Author contributions

WRB, JD, LBC and MMA designed the experiments. WRB performed the growth experiments in LBC’s lab. WB performed the initial sequencing data quality filtering, read mapping, identification of translated ORFs, correlations between proteomics and sequencing data. TT performed the differential gene expression and translational efficiency analyses as well as GO terms enrichment. SGM performed the subsampling analyses and correlations between different sets of sequencing data. BBM carried out the ribosome profiling protocol in JD’s lab. ACM performed the multidimensional scaling analysis. TT, WRB and MMA wrote the manuscript.

## Data availability

Supplementary data files have been uploaded to Figshare and can be accessed at http://dx.doi.org/10.6084/m9.figshare.5809812. This includes the transcriptome genomic coordinates, the gene table of counts, lisst of differentially expressed genes and gene/protein abundance estimates derived from RNA-Seq, Ribo-Seq and proteomics. The original sequencing data is at https://www.ncbi.nlm.nih.gov/bioproject/435567 (NCBI Bioproject PRJNA435567).

## Supplementary information

The supplementary file contains the supplementary tables and figures mentioned in the text.

## References

1. Rapaport, F. et al. Comprehensive evaluation of differential gene expression analysis methods for RNA-seq data. Genome Biol. 14, R95 (2013).

2. de Sousa Abreu, R., Penalva, L. O., Marcotte, E. M. & Vogel, C. Global signatures of protein and mRNA expression levels. Mol. Biosyst. 5, 1512–26 (2009).

3. Schwanhäusser, B. et al. Global quantification of mammalian gene expression control. Nature 473, 337–342 (2011).

4. Payne, S. H. The utility of protein and mRNA correlation. Trends Biochem. Sci. 40, 1–3 (2015).

5. Ponnala, L., Wang, Y., Sun, Q. & van Wijk, K. J. Correlation of mRNA and protein abundance in the developing maize leaf. Plant J. 78, 424–440 (2014).

6. Lee, M. V. et al. A dynamic model of proteome changes reveals new roles for transcript alteration in yeast. Mol. Syst. Biol. 7, 514–514 (2014).

7. Tebaldi, T. et al. Widespread uncoupling between transcriptome and translatome variations after a stimulus in mammalian cells. BMC Genomics 13, 220 (2012).

8. Ingolia, N. T., Ghaemmaghami, S., Newman, J. R. S. & Weissman, J. S. Genome-wide analysis in vivo of translation with nucleotide resolution using ribosome profiling. Science 324, 218–23 (2009).

9. Gerashchenko, M. V., Lobanov, A. V. & Gladyshev, V. N. Genome-wide ribosome profiling reveals complex translational regulation in response to oxidative stress. Proc. Natl. Acad. Sci. 109, 17394–17399 (2012).

10. Ji, Z., Song, R., Regev, A. & Struhl, K. Many lncRNAs, 5’UTRs, and pseudogenes are translated and some are likely to express functional proteins. Elife 4, e08890 (2015).

11. Mcmanus, C. J., May, G. E., Spealman, P. & Shteyman, A. Ribosome profiling reveals post-transcriptional buffering of divergent gene expression in yeast. 422–430 (2014).

12. Borg, I. & Groenen, P. J. F. Modern multidimensional scaling. (Springer, 1997).

13. Law, C. W., Chen, Y., Shi, W. & Smyth, G. K. voom: precision weights unlock linear model analysis tools for RNA-seq read counts. Genome Biol. 15, R29 (2014).

14. Morano, K. A., Grant, C. M. & Moye-Rowley, W. S. The response to heat shock and oxidative stress in Saccharomyces cerevisiae. Genetics 190, 1157–95 (2012).

15. Huang, D. W., Sherman, B. T. & Lempicki, R. A. Systematic and integrative analysis of large gene lists using DAVID bioinformatics resources. Nat. Protoc. 4, 44–57 (2009).

16. Zhong, Y. et al. RiboDiff: detecting changes of mRNA translation efficiency from ribosome footprints. Bioinformatics 33, 139–141 (2017).

17. Ingolia, N. T. Ribosome Footprint Profiling of Translation throughout the Genome. Cell 165, 22–33 (2016).

18. Jungfleisch, J. et al. A novel translational control mechanism involving RNA structures within coding sequences. Genome Res. 27, 95–106 (2017).

19. Yordanova, M. M. et al. AMD1 mRNA employs ribosome stalling as a mechanism for molecular memory formation. Nature 553, 356–360 (2018).

20. Aspden, J. L. et al. Extensive translation of small ORFs revealed by Poly-Ribo-Seq. Elife e03528 (2014).

21. Ruiz-Orera, J., Messeguer, X., Subirana, J. A. & Alba, M. M. Long non-coding RNAs as a source of new peptides. Elife 3, e03523 (2014).

22. Raj, A. et al. Thousands of novel translated open reading frames in humans inferred by ribosome footprint profiling. Elife 5, (2016).

23. Ruiz-Orera, J., Verdaguer-Grau, P., Villanueva-Cañas, J. L., Messeguer, X. & Albà, M. M. Translation of neutrally evolving peptides provides a basis for de novo gene evolution. Nat. Ecol. Evol. 2, 890–896 (2018).

24. Gerber, S. A., Rush, J., Stemman, O., Kirschner, M. W. & Gygi, S. P. Absolute quantification of proteins and phosphoproteins from cell lysates by tandem MS. Proc. Natl. Acad. Sci. 100, 6940–6945 (2003).

25. Edfors, F. et al. Gene-specific correlation of RNA and protein levels in human cells and tissues. Mol. Syst. Biol. 12, 883 (2016).

26. Slavoff, S. A. et al. Peptidomic discovery of short open reading frame-encoded peptides in human cells. Nat. Chem. Biol. 9, 59–64 (2013).

27. Stadler, M. & Fire, A. Conserved translatome remodeling in nematode species executing a shared developmental transition. Plos Genet. 9, e1003739 (2013).

28. Zid, B. M. & O’Shea, E. K. Promoter sequences direct cytoplasmic localization and translation of mRNAs during starvation in yeast. Nature 514, 117–121 (2014).

29. Khong, A. et al. The Stress Granule Transcriptome Reveals Principles of mRNA Accumulation in Stress Granules. Mol. Cell 68, 808–820.e5 (2017).

30. Luo, Y., Na, Z. & Slavoff, S. A. P-Bodies: Composition, Properties, and Functions. Biochemistry 57, 2424–2431 (2018).

31. Reid, D. W., Shenolikar, S. & Nicchitta, C. V. Simple and inexpensive ribosome profiling analysis of mRNA translation. Methods 91, 69–74 (2015).

32. Xie, S.-Q. et al. RPFdb: a database for genome wide information of translated mRNA generated from ribosome profiling. Nucleic Acids Res. 44, D254–D258 (2016).

33. Liu, W., Xiang, L., Zheng, T., Jin, J. & Zhang, G. TranslatomeDB: a comprehensive database and cloud-based analysis platform for translatome sequencing data. Nucleic Acids Res. 46, D206–D212 (2018).

34. Tsankov, A. M., Thompson, D. A., Socha, A., Regev, A. & Rando, O. J. The Role of Nucleosome Positioning in the Evolution of Gene Regulation. PLoS Biol. 8, e1000414 (2010).

35. Ingolia, N. T., Brar, G. a, Rouskin, S., McGeachy, A. M. & Weissman, J. S. The ribosome profiling strategy for monitoring translation in vivo by deep sequencing of ribosome-protected mRNA fragments. Nat. Protoc. 7, 1534–50 (2012).

36. Bolger, A. M., Lohse, M. & Usadel, B. Trimmomatic: a flexible trimmer for Illumina sequence data. Bioinformatics 30, 2114–20 (2014).

37. Martin, M. Cutadapt removes adapter sequences from high-throughput sequencing reads. EMBnet.journal 17.1, (2011).

38. Langmead, B., Trapnell, C., Pop, M. & Salzberg, S. L. Ultrafast and memory-efficient alignment of short DNA sequences to the human genome. Genome Biol. 10, R25 (2009).

39. Li, H. et al. The Sequence Alignment/Map format and SAMtools. Bioinformatics 25, 2078–2079 (2009).

40. Anders, S., Pyl, P. T. & Huber, W. HTSeq--a Python framework to work with high-throughput sequencing data. Bioinformatics 31, 166–9 (2015).

41. Robinson, M. D., McCarthy, D. J. & Smyth, G. K. edgeR: a Bioconductor package for differential expression analysis of digital gene expression data. Bioinformatics 26, 139–140 (2010).

42. Ashburner, M. et al. Gene ontology: tool for the unification of biology. The Gene Ontology Consortium. Nat. Genet. 25, 25–9 (2000).

