## Supplemental Tables and Figures for "Extensive post-transcriptional buffering of gene expression in the response to oxidative stress in baker’s yeast"

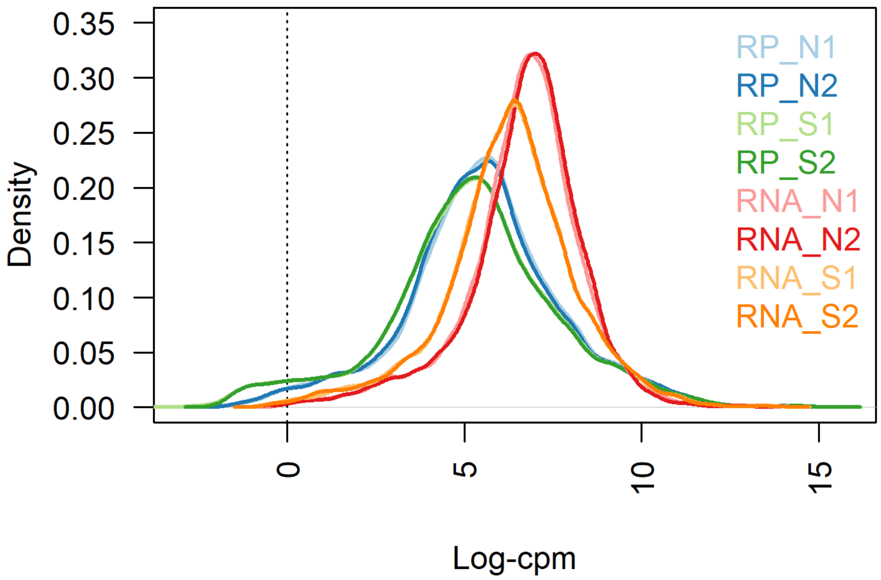


**Figure S1. Distribution of log2 CPM values for RNA-Seq and Ribo-Seq data.** RP: Ribo-Seq data; RNA: RNA-Seq data; N: normal growth conditions; S: stress conditions. Two sequencing replicates were generated per condition.

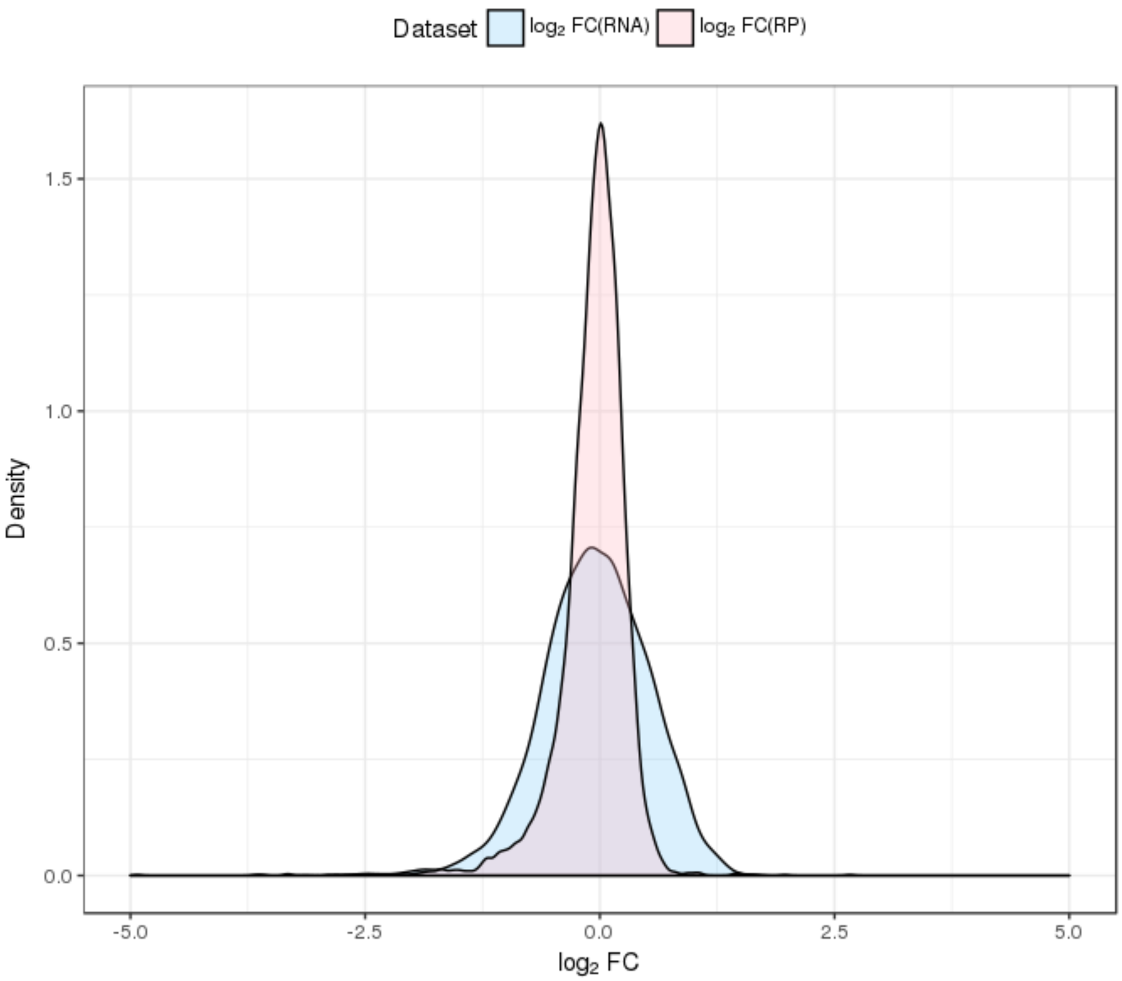


**Figure S2. Distribution of gene expression fold change (FC) differences in logarithmic scale using RNA-Seq and Ribo-Seq data.** The mapped reads were subsampled to have a similar number of reads for all samples. FC was calculated as the ratio between the number of reads in oxidative stress and normal conditions. We took the average number of reads per gene among the replicates of each experiment. The standard deviation of $\log_2$FC was 0.38 for Ribo-Seq (RP) and 0.56 for RNA-Seq (RNA).

### Tables

| | N1 | N2 | S1 | S2 |
|---|---|---|---|---|
| **RNA-Seq (RNA)** | 13,428,623 | 13,970,477 | 12,003,726 | 15,383,315 |
| **Ribo-Seq (RP)** | 36,295,110 | 34,464,847 | 31,796,926 | 31,207,820 |

**Table S1. Number of sequencing reads in different samples after the filtering steps.** N1 and N2 are normal growth conditions; S1 and S2 are oxidative stress conditions.

| | RNA_N1 | RNA_N2 | RNA_S1 | RNA_S2 | RP_N1 | RP_N2 | RP_S1 |
|---|---|---|---|---|---|---|---|
| RNA_N2 | 0.991 | 1 | | | | | |
| RNA_S1 | 0.948 | 0.926 | 1 | | | | |
| RNA_S2 | 0.942 | 0.918 | 0.997 | 1 | | | |
| RP_N1 | 0.843 | 0.802 | 0.868 | 0.867 | 1 | | |
| RP_N2 | 0.835 | 0.794 | 0.866 | 0.864 | 0.998 | 1 | |
| RP_S1 | 0.847 | 0.806 | 0.868 | 0.865 | 0.991 | 0.989 | 1 |
| RP_S2 | 0.839 | 0.798 | 0.865 | 0.863 | 0.992 | 0.991 | 0.999 |

**Table S2**. **Spearman correlation values for RNA-Seq and Ribo-Seq.** Gene expression units are log2CPM, where CPM is counts per million. RP: Ribo-Seq. RNA: RNA-Seq. N1: normal, replicate 2; N2: normal, replicate 2; S1: stress, replicate 1; S2: stress, replicate 2.

| | N1 | N2 | N3 | S1 | S2 |
|---|---|---|---|---|---|
| N2 | 0.909 | 1 | | | |
| N3 | 0.928 | 0.908 | 1 | | |
| S1 | 0.93 | 0.884 | 0.913 | 1 | |
| S2 | 0.904 | 0.923 | 0.9 | 0.899 | 1 |
| S3 | 0.856 | 0.863 | 0.882 | 0.829 | 0.858 |

**Table S3. Spearman correlation values for proteomics data.** Protein expression units are nomalized (equalization of the medians) area, considering only proteins for which at least 3 unique peptides were detected- 2200 genes in total. N1: normal, replicate 2; N2: normal, replicate 2; N3: normal, replicate 3; S1: stress, replicate 1; S2: stress, replicate 2; S3: stress, replicate 3.

| Gene_name | Genomic_coord_CDS | Prot_normal | Coord_norm | Prot_stress | Coord_stress |
|---|---|---|---|---|---|
| CRF1 | NC_001136.10:912099-913502 | rna1169-1 | 912099-913502 | rna1169-3 | 912448-912504 |
| YNL046W | NC_001146.8:542304-542822 | rna5108-2 | 542365-542502 | rna5108-1 | 542304-542822 |
| YKR075C | NC_001143.9:579827-580750 | rna3731-7 | 580108-580131 | rna3731-1 | 579827-580750 |
| YJL118W | NC_001142.9:191639-192298 | rna3138-3 | 191647-191688 | rna3138-1 | 191639-192298 |
| MMS22 | NC_001144.5:771940-776304 | rna4176-6 | 771992-772102 | rna4176-1 | 771940-776304 |
| TOS8 | NC_001139.9:325331-326161 | rna2107-3 | 326070-326150 | rna2107-1 | 325331-326161 |
| MET16 | NC_001148.4:876847-877632 | rna6262-2 | 876861-877034 | rna6262-1 | 876847-877632 |

**Table S4. Alternative translated ORFs.** List of genes for which the longest ORF with translation evidence was different for normal and stress conditions. Translation evidence was predicted with RibORF using a score cut-off of 0.7. Genomic_coord_CDS: chromosome name and position of the annotated coding sequence (CDS). Only cases in which the annotated CDS was covered by more than 200 Ribo-Seq reads in at least one of the samples were considered.

| | N1 | N2 | S1 | S2 |
|---|---|---|---|---|
| **RNA-seq (RNA)** | 15,334,498 | 15,302,667 | 15,353,235 | 15,341,682 |
| **Ribo-Seq (RP)** | 15,515,017 | 15,387,974 | 15,482,027 | 15,248,146 |

**Table S5. Total number of counts after subsampling.** Subsampling was performed taking a random set of reads from each sample so that the different samples had similar number of counts.

| | RP_N1 | RP_N2 | RP_S1 | RP_S2 | RNA_N1 | RNA_N2 | RNA_S1 | RNA_S2 |
|---|---|---|---|---|---|---|---|---|
| RP_N1 | 1 | | | | | | | |
| RP_N2 | 0.99 | 1 | | | | | | |
| RP_S1 | 0.99 | 0.99 | 1 | | | | | |
| RP_S2 | 0.99 | 0.99 | 0.99 | 1 | | | | |
| RNA_N1 | 0.88 | 0.88 | 0.87 | 0.87 | 1 | | | |
| RNA_N2 | 0.84 | 0.84 | 0.84 | 0.83 | 0.99 | 1 | | |
| RNA_S1 | 0.88 | 0.88 | 0.87 | 0.87 | 0.95 | 0.93 | 1 | |
| RNA_S2 | 0.88 | 0.88 | 0.87 | 0.87 | 0.94 | 0.92 | 0.99 | 1 |

**Table S6. Spearman correlation values for RNA-Seq and Ribo-Sseq after subsampling.** Gene expression units are log2CPM, where CPM is counts per million.

| Gene Name | Description |
|---|---|
| DIB1 | 17-kDa component of the U4/U6aU5 tri-snRNP; plays an essential role in pre-mRNA splicing; human ortholog TXNL4A (the human U5-specific 15-kDa protein) complements yeast dib1 null mutant |
| SPC19 | Essential subunit of the Dam1 complex (aka DASH complex); complex couples kinetochores to the force produced by MT depolymerization thereby aiding in chromosome segregation; also localized to nuclear side of spindle pole body |
| BNS1 | Protein of unknown function; overexpression bypasses need for Spo12p, but not required for meiosis; BNS1 has a paralog, SPO12, that arose from the whole genome duplication |
| YOR019W | Protein of unknown function; may interact with ribosomes, based on co-purification experiments; YOR019W has a paralog, JIP4, that arose from the whole genome duplication |
| RMI1 | Subunit of the RecQ (Sgs1p) - Topo III (Top3p) complex; stimulates superhelical relaxing, DNA catenation/decatenation and ssDNA binding activities of Top3p; involved in response to DNA damage; functions in S phase-mediated cohesion establishment via a pathway involving the Ctf18-RFC complex and Mrc1p; stimulates Top3p DNA catenation/decatenation activity; null mutants display increased rates of recombination and delayed S phase |
| HTL1 | Component of the RSC chromatin remodeling complex; RSC functions in transcriptional regulation and elongation, chromosome stability, and establishing sister chromatid cohesion; involved in telomere maintenance |
| NDT80 | Meiosis-specific transcription factor; required for exit from pachytene and for full meiotic recombination; activates middle sporulation genes; competes with Sum1p for binding to promoters containing middle sporulation elements (MSE) |
| BUB3 | Kinetochore checkpoint WD40 repeat protein; localizes to kinetochores during prophase and metaphase, delays anaphase in the presence of unattached kinetochores; forms complexes with Mad1p-Bub1p and with Cdc20p, binds Mad2p and Mad3p; functions at kinetochore to activate APC/C-Cdc20p for normal mitotic progression |
| IML3 | Outer kinetochore protein and component of the Ctf19 complex; involved in the establishment of pericentromeric cohesion during mitosis; prevents non-disjunction of sister chromatids during meiosis II; forms a stable complex with Chl4p; required for localization of Sgo1p to pericentric sites during meiosis I; orthologous to human centromere constitutive-associated network (CCAN) subunit CENP-L and fission yeast fta1 |
| DMC1 | Meiosis-specific recombinase required for double-strand break repair; also required for pairing between homologous chromosomes and for the normal morphogenesis of synaptonemal complex; binds ssDNA and dsDNA, forms helical filaments; potent inhibitor of the ATPase activity of Srs2p helicase, blocking its ssDNA translocating motor activity and inhibiting its antirecombinase activity; stimulated by Rdh54p; homolog of Rad51p and the bacterial RecA protein |
| CDC26 | Subunit of the Anaphase-Promoting Complex/Cyclosome (APC/C); which is a ubiquitin-protein ligase required for degradation of anaphase inhibitors, including mitotic cyclins, during the metaphase/anaphase transition; relocalizes to the cytosol in response to hypoxia |
| SDC25 | Non-essential Ras guanine nucleotide exchange factor (GEF); localized to the membrane; expressed in poor nutrient conditions and on nonfermentable carbon sources; contains a stop codon in S288C, full-length gene includes YLL017W; SDC25 has a paralog, CDC25, that arose from the whole genome duplication |

**Table S7. List of putatively repressed genes related to the cell cycle.** These genes showed significantly decreased translational efficiency during stress and were significantly under-expressed during stress using Ribo-Seq data, suggesting their translation is inhibited when compared to the rest of genes.
